# BMP signaling orchestrates a transcriptional network to control the fate of mesenchymal stem cells (MSCs)

**DOI:** 10.1101/104927

**Authors:** Jifan Feng, Junjun Jing, Jingyuan Li, Hu Zhao, Vasu Punj, Tingwei Zhang, Jian Xu, Yang Chai

## Abstract

Signaling pathways are used reiteratively in different developmental processes yet produce distinct cell fates through activating specific downstream transcription factors. In this study, we used tooth root development as a model to investigate how the BMP signaling pathway regulates specific downstream transcriptional complexes to direct the fate determination of multipotent mesenchymal stem cells (MSCs). We first identified the MSC population supporting mouse molar root growth as Gli1+ cells. Using a Gli1-mediated transgenic animal model, our results provide the first *in vivo* evidence that BMP signaling activity is required for the odontogenic differentiation of MSCs. Specifically, we identified transcription factors that are downstream of BMP signaling and are expressed in a spatially restricted pattern consistent with their potential involvement in determining distinct cellular identities within the dental mesenchyme. Finally, we found that overactivation of one key transcription factor, Klf4, associated with the odontogenic region, promotes odontogenic differentiation of MSCs. Collectively, our results demonstrate the functional significance of BMP signaling in regulating the fate of MSCs during root development and shed light on how BMP signaling can achieve functional specificity in regulating diverse organ development.

**Summary Statement:** BMP signaling activity is required for the lineage commitment of MSCs and transcription factors downstream of BMP signaling may determine distinct cellular identities within the dental mesenchyme.

## INTRODUCTION

During development and throughout life, tissue growth and homeostasis require tightly regulated proliferation and differentiation of immature precursor cells. Multipotent mesenchymal stem cells (MSCs), first reported in bone marrow tissues, have been shown in a variety of mesenchymal tissues with different developmental origins and physiological functions (Friedenstein et al., 1968; Friedenstein et al., 1976; Bianco et al., 2008). These MSCs from different tissues are identified based on their common *in vitro* defining characteristics including colony-forming ability, multipotency (osteo-, chondro-and adipogenic potentials), and the expression of MSC surface markers (Dominici et al., 2006). Despite these *in vitro* similarities, MSCs from various tissues undergo strict lineage-specific differentiation programs *in vivo*, faithful to their unique tissue origins and environment (Gronthos et al., 2000; Beederman et al., 2013). To date, the ontology and niche environment required for the MSCs supporting specific tissue growth have yet to be clearly elucidated.

During embryonic development, multipotent cells with MSC characteristics have been identified in sites where post-migratory cranial neural crest cells (CNCCs) reside (Chung et al., 2009). Mouse molar development includes crown formation preceding root initiation, followed by root elongation. Molars cease growing after their development and thus represent a good model for studying human tooth development. During late stages of molar development, the molar epithelium tissue degenerates and dissociates, due to the loss of Sox2+ epithelial stem cells regulated by a BMP-SHH signaling cascade (Juuri et al., 2012; Li et al., 2015). In contrast, root development occurs mainly in the CNCC-derived root mesenchymal tissue that forms the future pulp, dentin and periodontium.

This restricted apical growth of the molar also coincides with the presence of a distinct MSC population in humans, namely stem cells of the apical papilla (SCAPs) (Sonoyama et al., 2006; Sonoyama et al., 2008). SCAPs exhibit classical MSC characteristics, as well as higher colony forming efficiency and growth potential capacity compared with stem cells from the dental pulp of the adult tooth (Sonoyama et al., 2006; Sonoyama et al., 2008). However, it remains unclear whether mice have a similar MSC counterpart to human SCAPs and how this stem cell population undergoes odontogenic lineage commitment *in vivo* during tooth morphogenesis. The only MSCs that have been definitively characterized reside in adult mouse incisors that grow continuously throughout life (Zhao et al., 2014). Although recent studies have shown that blood vessel walls harbor a reservoir of MSCs in multiple tissues (Shi and Gronthos, 2003; Crisan et al., 2008), our cell lineage tracing experiments have shown that the stem cell population supporting tissue growth *in vivo* is supported by Gli1+ cells, the majority of which are not perivascular cells in adult mouse incisors or are not associated with the vasculature in the cranial sutures (Zhao et al., 2014; Zhao et al., 2015). In addition, lineage tracing of NG2+ perivascular cells in mouse molars demonstrated that they exhibit only a limited contribution to molar mesenchyme development (Feng et al., 2011; Zhao et al., 2014), so the identity and location of mouse molar MSCs remains unclear.

Cell fate determination during tissue patterning and lineage commitment is often coordinated by signaling pathways via their activation or inhibition of transcription factors. These transcription factors regulate gene expression by acting synergistically or antagonistically, forming gene regulatory networks. During early embryonic development, BMPs play a critical role in fate determination including regulating cell fate, growth and patterning (Helms et al., 1998; Zhang et al., 2013). Similarly, in the craniofacial region, mandibular domain identities are determined by the mutually antagonistic relationship of Bmp and Fgf signals from the oral epithelium differentially regulating transcription factors Msx1/2, Barx1, Dlx2 and Lhx6/7 in the CNC-derived mesenchyme (Tucker and Sharpe, 2004). In adult organs, BMP signaling helps maintain the stem cell population size by inhibiting stem cell proliferation and niche expansion (He et al., 2004). BMP signaling activity is also required for activating MSCs to undergo differentiation in cell culture (Beederman et al., 2013; Wei et al., 2013). However, it is not clear how BMP signaling controls MSC cell fate determination during tooth root development.

Here, we have identified the precise *in vivo* identity of the MSC population critical for postnatal tooth development in the apical region of molars. These Gli1+ cells are adjacent to, but more apical than, cells with active BMP signaling. Our results indicate that BMP signaling activity is indispensable for the activation of MSCs into their differentiation program. In addition, we identified potential downstream targets of BMP signaling, including spatially restricted transcription factors that may activate the odontogenic mesenchyme lineage program, such as Klf4 and Lhx8. Thus, our findings suggest that BMP signaling orchestrates a transcriptional network that regulates mesenchymal stem cell lineage commitment.

## RESULTS

### Identification of putative mesenchymal stem cells (MSCs) in the developing molar apical mesenchyme

Gli1 was recently identified as an *in vivo* marker for adult mouse incisor MSCs (Zhao et al., 2014), but it remained unclear if a similar population of stem cells transiently exists to support mouse molar development. Our recent study also provided in vivo evidence that Hh signaling participates in root development and that Gli1+ cells are critical for root formation (Liu et al., 2015). To detect putative molar MSCs, we examined the detailed Gli1 expression pattern in the molar apical mesenchyme during root formation. At birth, Gli1 was expressed throughout the apical half of the dental mesenchyme (Fig. 1A). From this stage onwards, Gli1+ cells became gradually restricted to the most apical region (Fig. 1A-D) and eventually was undetectable by the adult stage (Zhao et al., 2014), consistent with a transient presence of stem cells during root development. We performed lineage tracing of Gli1+ cells from postnatal day 3.5 (P3.5), just prior to root formation initiation (at P7.5). One day after labeling, Gli1+ cells (tdTomato+) were located at the most apical region of the dental mesenchyme (Fig. 1E-F), with comparable expression to *Gli1-LacZ* mice. After two weeks, the progeny of these Gli1+ cells were detectable throughout the newly formed root odontoblasts and pulp cells (Fig. 1G-H), indicating that the progeny of these Gli1+ cells contribute to the entire root mesenchyme. Furthermore, *in vitro* assays showed that Gli1+ cells also have classic MSC characteristics including colony formation (Fig. 1I) and multi-lineage differentiation into osteoblasts, chondrocytes and adipocytes (Fig. 1J-L). Taken together, these results indicate that these Gli1+ cells function as *in vivo* MSCs that support postnatal molar mesenchymal development.

**Figure 1.**
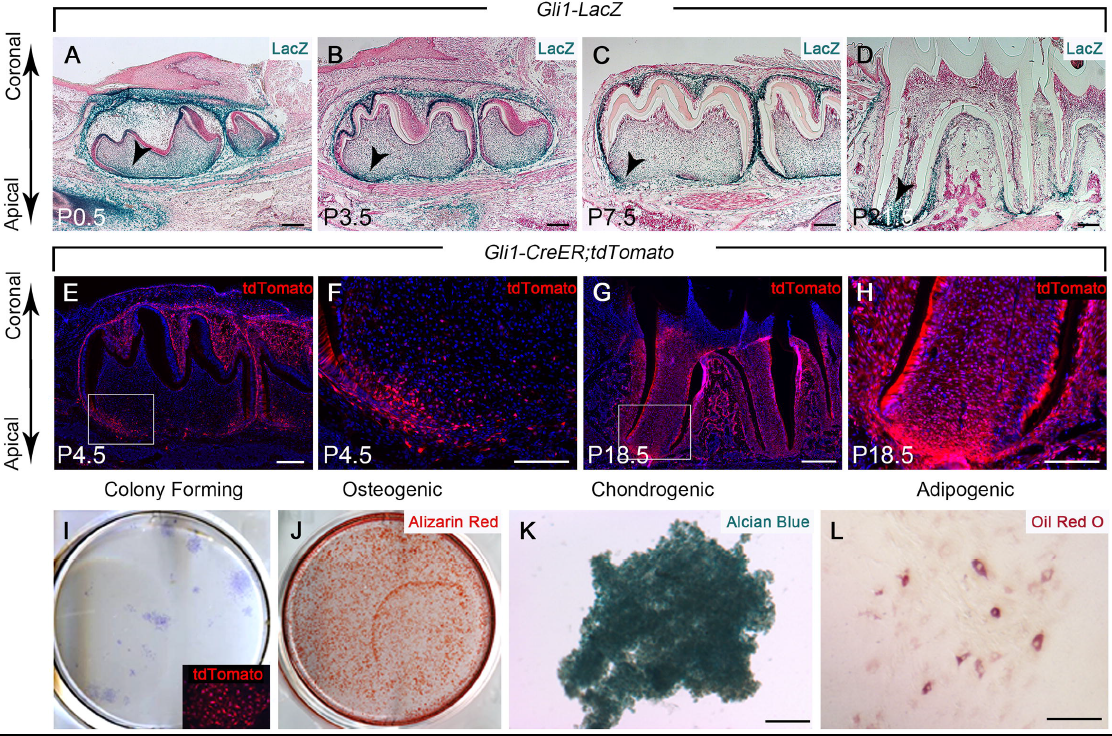
Gli1 is an in vivo marker for MSCs in the developing molar apical mesenchyme. (A-D) LacZ staining (blue) of sagittal sections of mandibular molars from *Gli1-LacZ* mice at P0.5, P3.5, P7.5, and P21.5. Arrowheads indicate Gli1+ cells in the apical region of the mesenchyme. (E-H) Visualization of sagittal sections of mandibular molars from *Gli1-CreER;tdTomato* mice at P4.5 and P18.5 after induction at P3.5. The progeny of the Gli1 lineage appear red. Boxes in E and G are shown magnified in F and H, respectively. (I-L) Colony forming assay and osteogenic, chondrogenic, and adipogenic differentiation assays of cells from the Gli1+ region in the apical mesenchyme of molars from P5.5 *Gli1-CreER;tdTomato* mice induced at P3.5. Toluidine blue staining was used to visualize colony formation after culture for two weeks (I). Insert shows colonies are derived from Gli1+ cells (red). Alizarin Red (J), Alcian Blue (K), and Oil Red O (L) staining to detect osteogenic, chondrogenic, and adipogenic differentiation after three weeks. Scale bars, 100μm.

### Activated BMP signaling in the dental mesenchyme as these cells become committed to differentiate into odontoblasts

Because tooth development occurs from the crown to the root, cells in more coronal regions are progressively more differentiated than those in the apical region, consistent with the presence of Gli1+ MSCs at the most apical part of the postnatal developing molar mesenchyme. Previous studies have reported the presence of Bmp ligands during all stages of tooth development, but the pattern of active BMP signaling remained unknown. We analyzed activated BMP signaling during root formation using pSmad1/5/8 expression as a readout. At E18.5, prior to the initiation of molar root formation, BMP activity is widespread but excluded from the apical region (Fig. S1A). From this stage onwards, BMP signaling persists in the developing molar mesenchyme, whereas the zone of exclusion becomes increasingly restricted in the apical region (Fig. S1A-D). In contrast, the zone of exclusion for BMP activity in the incisor is maintained postnatally and throughout adulthood (Fig. S1E-H). This difference is consistent with the persistence of stem cells in the incisor and the gradual loss of stem cells in the molar during later development. To investigate the relationship between BMP signaling and Gli1+ MSCs at the onset of root initiation, we examined their expression pattern at P3.5. Activated BMP signaling (pSmad1/5/8) was localized more coronally than the Gli1+ cells in the most apical region (Fig. 2A-C), suggesting that active BMP signaling is associated with more committed cells adjacent to but excluded from the Gli1+ MSCs. Similarly, lineage tracing of Gli1+ cells confirmed this absence of BMP activity in the Gli1+ MSC region (Fig. 2D). Two days after induction, Gli1+ cells were only present in the most apical region (Fig. 2D”) and BMP signaling was more coronal than the Gli1+ cells (Fig. 2D’). At later time points, progeny of these Gli1+ cells became more committed as odontoblasts and other dental pulp cells, and they co-localized to the region with activated BMP signaling (Fig. 2E), consistent with a requirement for BMP signaling for MSCs to become committed as odontoblasts and pulp tissue.

**Figure 2.**
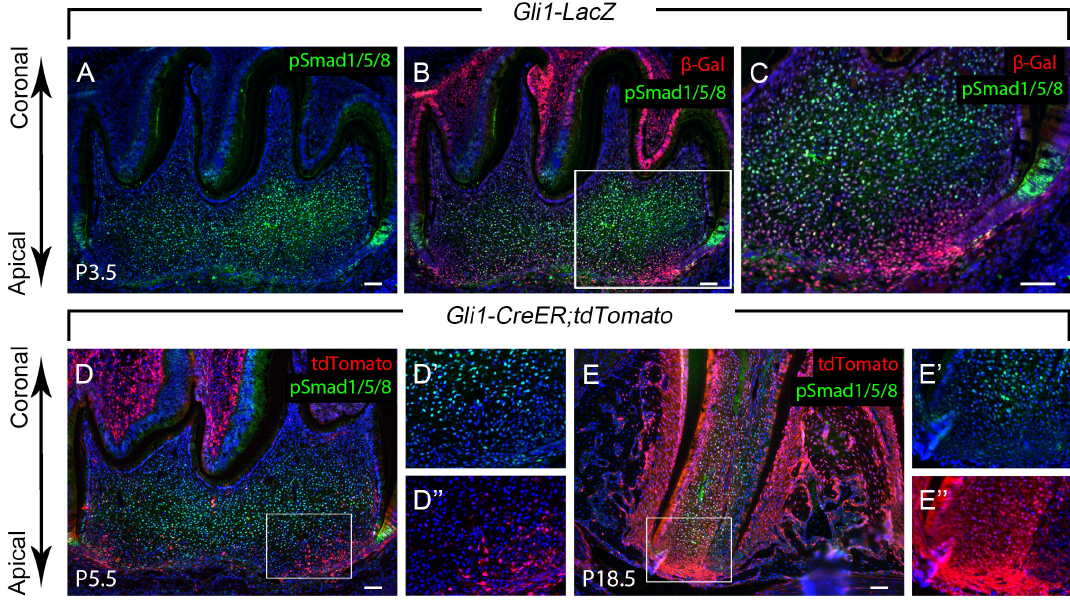
Co-localization of activated BMP signaling (pSmad1/5/8) and Gli1» MSCs and their progeny in developing roots. (A-C) pSmad1/5/8 (green) andβ-gal (red) immunostainingof sagittal sections of mandibular molars from P3.5 *Gli1-LacZ* mice. pSmad1/5/8 indicates activated BMP signaling and β-gal indicates Gli1 expression. Box in B is shown magnified in C. (D-E) pSmad1/5/8 immunostaining (green) and visualization of tdTomato (red) of sagittal sections of mandibular molars from *Gli1-CreER;tdTomato* mice at P5.5 and P18.5 after induction at P3.5. The progeny of the Gli1 lineage appear red. Boxes in D and E are shown magnified in D’ and D” and E’ and E”, respectively. D’ and E’ show pSmad1/5/8 staining and D” and E” show tdTomato visualization alone. Scale bars, 100μm.

### BMP signaling is indispensable for Gli1+ MSCs to initiate the apical growth of the root

Disruption of BMP signaling in the root mesenchyme before root formation in *Gli1-CreER;Bmpr1*α^*fl/fl*^ mice resulted in impaired root development (Fig. 3A-B, E-F). At P18.5, roots were well developed in control mice and the tooth was ready to erupt (Fig. 3A,B), whereas there was no histological structure resembling a root in *Gli1-CreER;Bmpr1*α^*fl/fl*^ mice, although crown and root patterning appeared to be unaffected (Fig. 3E,F). In addition, dentin formation was defective and no cells showing typical columnar odontoblast morphology were detectable in *Gli1-CreER;Bmpr1*α^*fl/fl*^ mice (Fig. 3E,F). Consistent with this, we failed to detect expression of odontoblast differentiation marker *Dspp* (dentin sialophosphoprotein) in the most apical region of *Gli1-CreER;Bmpr1*α^*fl/fl*^ mouse molars (Fig. 3C,G), suggesting there is a functional requirement for BMP signaling to initiate odontogenic differentiation in Gli1+ MSCs. Next, we performed lineage tracing of Gli1+ cells after loss of BMP signaling and found that their derivatives accumulated in the periapical region (Fig. 3H), failing to grow apically as in control mice (Fig. 3D). Moreover, the region of proliferative cells in the apical mesenchyme was expanded in *Gli1-CreER;Bmpr1*α^*fl/fl*^ mice at P7.5, particularly in the region close to the preodontoblast/odontoblast cells (Fig. 3I-L), possibly due to the failure of these cells to enter into the odontogenic differentiation program. In contrast, ablating BMP signaling in the dental epithelium using *K14-rtTA;tetO-Cre;Bmpr1*α^*fl/fl*^ mice showed that epithelial BMP signaling is not specifically required to regulate the root pattern, length or dentin formation at postnatal stages (Fig. S2). Therefore, although Gli1+ cells are also present in the dental epithelial cells, BMP signaling is specifically required to regulate the differentiation of MSCs during postnatal tooth root formation.

**Figure 3.**
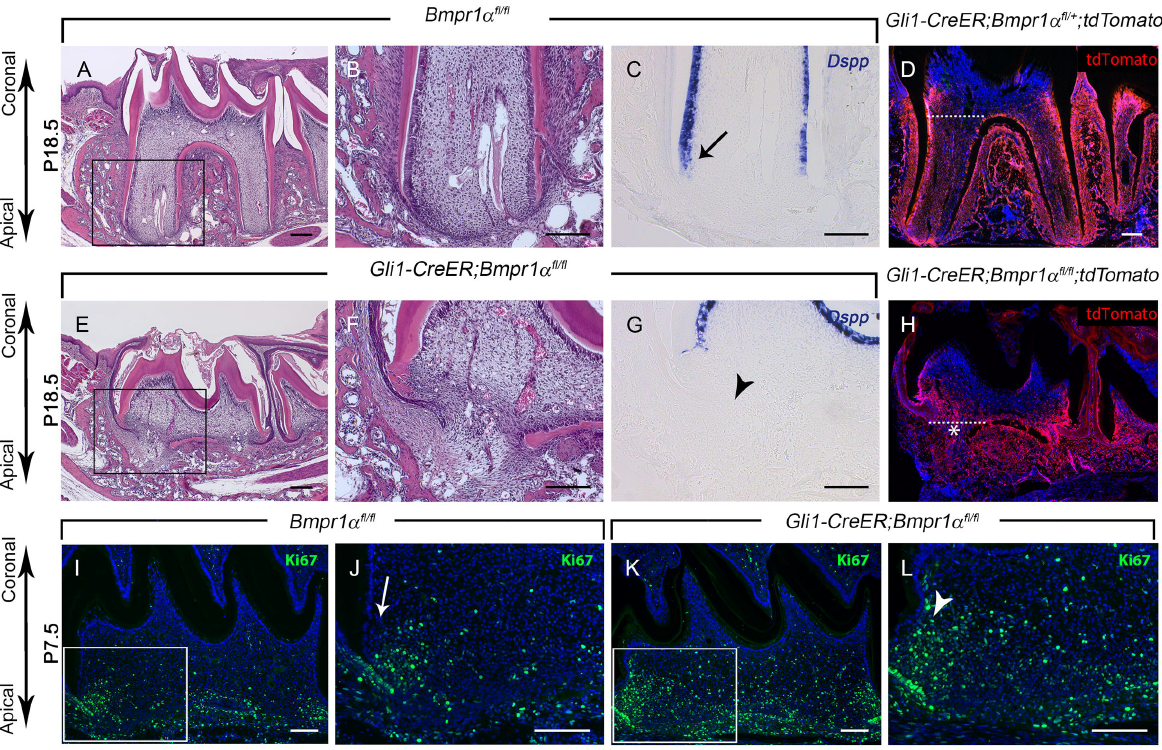
Disruption of BMP signaling in the root mesenchyme results in a differentiation defect. (A-C, E-G) H&E staining (A-B, E-F) and*Dspp*in situ hybridization (purple; C,G) of sagittal sections of mandibular molars of E18.5 littermate *Bmpr1*α^*fl/fl*^ control and *Gli1-CreER;Bmpr1*α^*fl/fl*^ mice after induction at P3.5. Boxes in A and E are shown magnified in B andF, respectively. Arrow indicates positive *Dspp* staining (C) and arrowhead indicates absence of staining (G). (D, H) Visualization of sagittal sections of mandibular molars from P18.5 littermate *Gli1-CreER;Bmpr1*α^*fl/+*^; *tdTomato* control and *Gli1-CreER;Bmpr1*α^*fl/fl*^; *tdTomato* mice after induction at P3.5. The progeny of the Gli1 lineage appear red. Asterisk indicates Gli1 derivatives in the periapical region and dotted lines indicate crown-root junction. (I-L) Ki67 immunostaining (green) of P7.5 littermate *Bmpr1* α^*fl/fl*^ control and *Gli1-CreER;Bmpr1*α^*fl/fl*^ mice induced P3.5. Boxes in I and K are shown magnified in J and L, respectively. Arrowhead indicates the expansion of the Ki67+ proliferative area (L) compared to control (arrow in J). Scale bars, 100μm.

### BMP signaling orchestrates a transcriptional network regulating mesenchymal stem cell lineage commitment

To identify downstream targets of BMP signaling in Gli1+ MSCs as they become committed into odontoblasts, we analyzed gene expression profiles in the apical half of the molar mesenchyme using RNA-seq from *Gli1-CreER;Bmpr1*α^*fl/fl*^ and control mice at P7.5. We harvested samples four days after tamoxifen induction because it takes approximately two days for tamoxifen to mediate Cre recombination and two days for Gli1+ MSCs to start to commit into pre-odontoblasts, based on our results showing that proliferation was altered using the same induction regimen (see Fig. 3I-L). We found 405 genes with altered expression in *Gli1-CreER;Bmpr1*α^*fl/fl*^ samples, using the criteria of >1.5 fold change and p-value < 0.05. Of these, 269 were upregulated and 136 were downregulated (accession number GSE79791), as indicated by representative mouse genome screen shots (Fig. 4A). Pathway enrichment assay demonstrated that, in addition to the BMP2 target genes, extracellular matrix proteins, cell adhesion, facial morphology, receptor binding, and cell proliferation related genes were all amongst the most enriched categories (Fig. 4B). We identified 29 transcription factors from this group, because these regulatory genes play critical roles in cell fate determination (Table S1). In particular, we focused on transcription factors associated with early craniofacial and tooth development, such as Klf4. In addition, we identified other factors such as Pax9 as interesting candidates among the altered genes.

**Figure 4.**
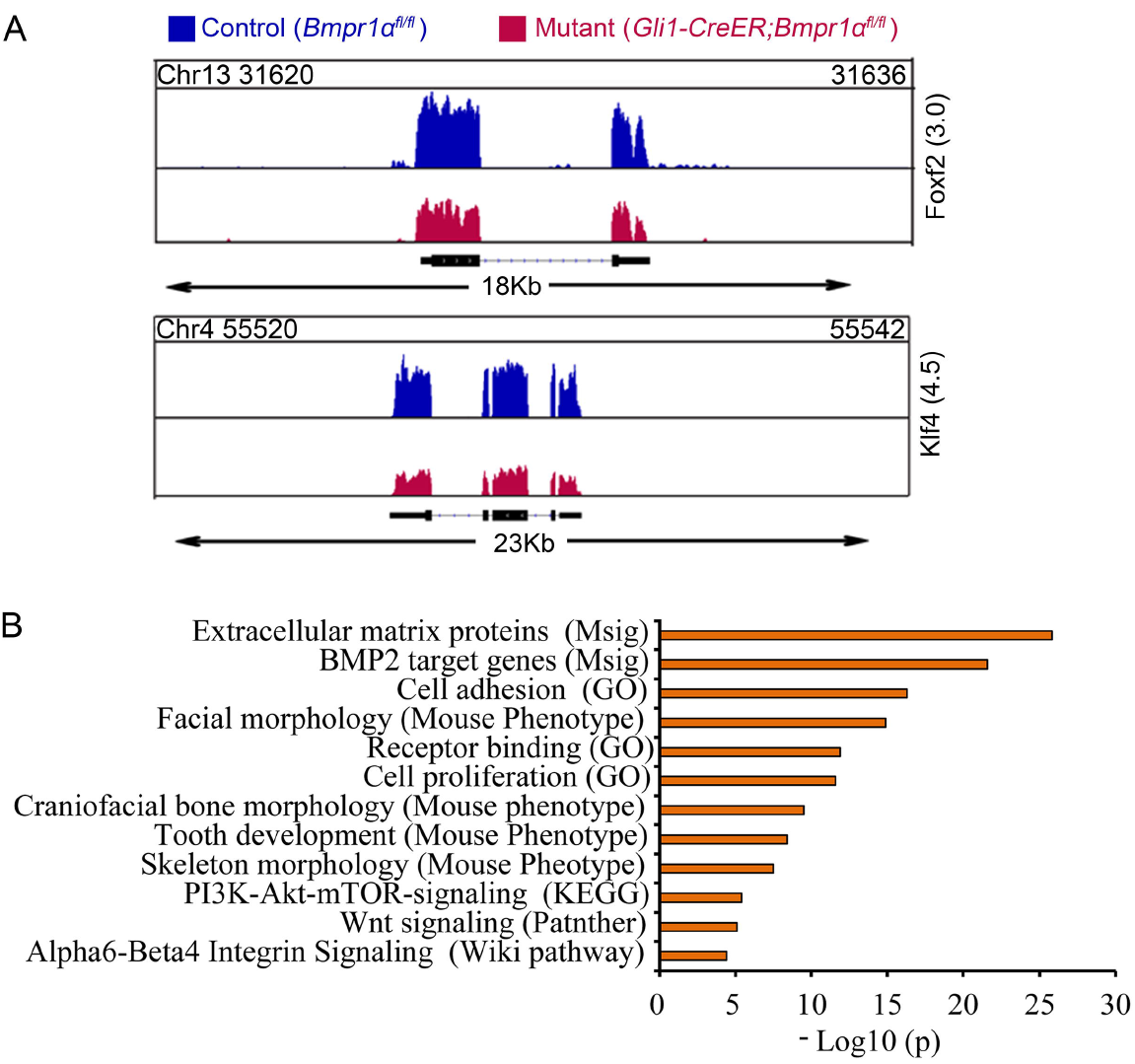
Downstream target gene analysis of *Gli1-CreER;Bmpr1*α^*fl/fl*^ mutant mice compared with*Bmpr1*α^*fl/fl*^ control mice. (A) Mouse genomic snapshots of two representative genes (Klf4 and Foxf2) identified in RNA-seq analysis of *Gli1-CreER;Bmpr1*α^*fl/fl*^ apical molars, showing downregulation (blue) relative to their matched controls (red). Genomic location of each gene is shown below. Numbers in parentheses on the right indicate sequencing depth. (B) Pathway analysis from RNA-seq data. Each enriched pathway is ranked based on p-value that was computed from the biochemical distribution and independence for probability. The database is indicated in the parentheses.

Because the RNA-seq experiment assayed a heterogeneous population, we confirmed the alteration in expression of interesting candidates in the apical mesenchyme *in vivo*. At P7.5, Gli1+ cells have begun to contribute to a small amount of growth resulting in a protrusion of the apical papilla (see Fig. 1C), whereas BMP activity was located more coronally in the pulp as well as in the pre-odontoblast/odontoblast region (Fig. 5A). In *Gli1-CreER;Bmpr1*α^*fl/fl*^ mice, the region of pSmad1/5/8 activity has been restricted more coronally and appeared undetectable in the pre-odontoblast/odontoblast region and in the pulp mesenchyme (Fig. 5B). In the same region, we observed several spatially restricted patterns of transcription factors, suggesting their involvement in determining distinct cellular identities within the dental mesenchyme. For example, in control mice, transcription factors such as Pax9 were detectable in the most apical region (Fig. 5C), comparable to Gli1 expression. In contrast, Fox2 was detectable in the pulp mesenchyme excluding the odontoblast layer (Fig. 5E), whereas Lhx8, Satb2 and Klf4 were closely associated with the odontogenic (pre-odontoblast/odontoblast) region (Fig. 5G, I, K). Furthermore, we found that loss of BMP signaling in the apical region leads to alterations in this network. In *Gli1-CreER;Bmpr1*α^*fl/fl*^ mice, Pax9 expression was expanded more coronally towards the pulp and odontogenic region (Fig. 5D), Foxf2, Lhx8 and Satb2 were reduced (Fig. 5F, H, J), and Klf4 expression was undetectable (Fig. 5L). These data suggest that BMP signaling fine-tunes the spatial distribution of this signaling network by activating differentiation related transcription factors and inhibiting transcription factors involved in stem cell maintenance.

**Figure 5.**
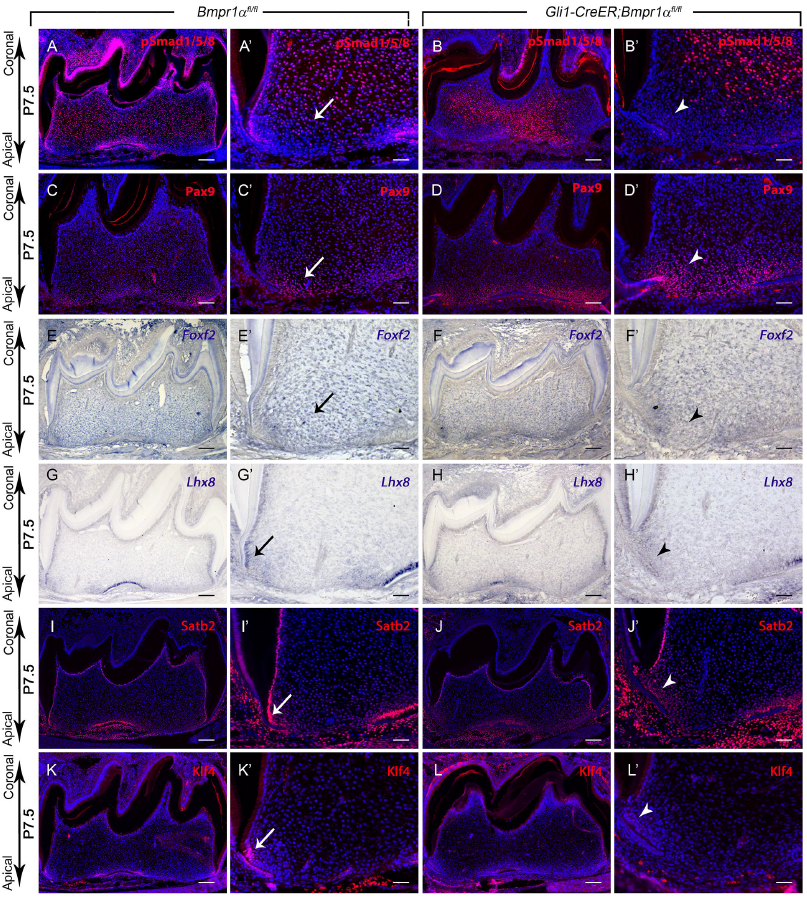
Altered spatial expression patterns of putative downstream factors after loss of BMP signaling in the molar apical mesenchyme. Immunostaining (red) of pSmad1/5/8 (A,B), Pax9 (C,D), Satb2 (I,J), and Klf4 (K,L) and in situ hybridization of *Foxf2* (E,F) and *Lhx8* (G,H) of P7.5 littermate *Bmpr1*α^*fl/fl*^ control and *Gli1-CreER;Bmpr1* α^*fl/fl*^ mice induced P3.5. A’-L’ are magnified images from A-L, respectively. Arrows indicate positive staining in control samples. Arrowheads indicate altered staining in targeted region of mutant samples. Scale bars: A-L, 100μm; A’-L’, 50μm.

### BMP regulates dental pulp tissue differentiation via activation of Klf4 expression

Amongst the group of transcription factors associated with pre-odontoblast/odontoblast identity, we noticed that the expression of Klf4 was closely adjacent to but excluded from the proliferative cells (Fig. 6A,A’, B,B’), suggesting that Klf4 may promote cell differentiation. Moreover, we found that Klf4 expression was restricted to the pre-odontoblast region and appeared decreased in the region of mature odontoblasts (Fig. 6A), suggesting that Klf4 might function as a switch for odontoblast differentiation. We showed above that pSmad1/5/8 overlapped with Klf4 expression in the apical region of the molar mesenchyme (see Fig. 5A, A’, K, K’). In addition, Bmp2 or Bmp4 treatment resulted in elevated Klf4 expression in dental pulp cells (Fig. 6C). Furthermore, consistent with a recent study that reported that Smad1/5 complex binds to the promoter region of Klf4 in mESCs (mouse embryonic stem cells) (Morikawa et al., 2016), we found that Klf4 also co-immunoprecipitates with Smad1 (Fig. 6D), suggesting that Klf4 is a direct target of BMP signaling. To test whether Klf4 may play a key role in regulating odontogenic differentiation, we used an adenovirus vector approach to activate it transiently in the apical mesenchyme of control P5.5 molar explants *in vitro*. We found that mouse Klf4 proteins were robustly expressed following infection with adenovirus Ad-m-Klf4 vectors (Fig. 6E, E’). Moreover, overexpression of Kfl4 promoted odontoblast differentiation based on its significant activation of odontoblast-specific marker *Dspp* in dissociated apical pulp culture (Fig. 6F). Taken together, our data suggests that Klf4 is spatially restricted to the odontogenic region and may be a key regulator of the transcriptional network downstream of BMP that controls odontoblast differentiation of MSCs during root development.

**Figure 6.**
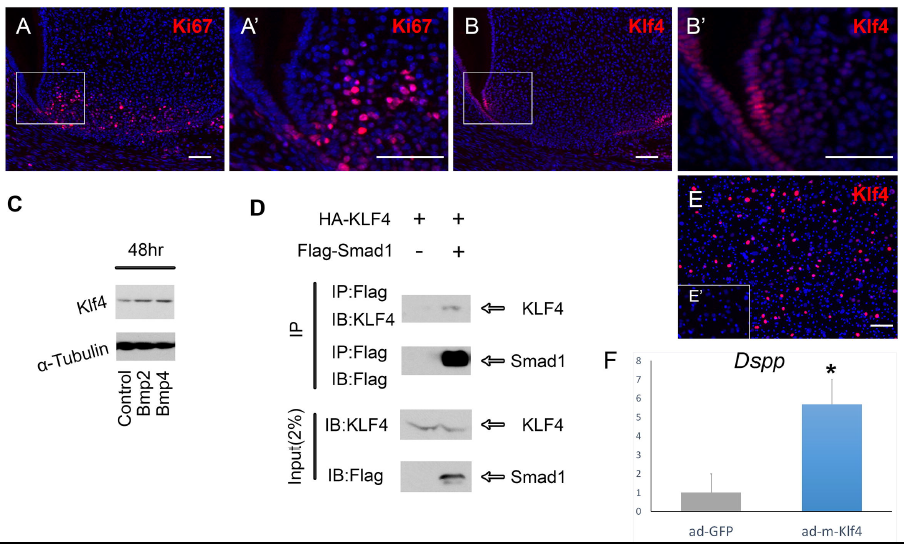
Activation of Klf4 in dental apical mesenchyme explants activates expression of odontoblast marker *Dspp*. (A-B) Ki67 and Klf4 immunostaining (red) of molars from P7.5 *Gli1-CreER;Bmpr1*α^*fl/fl*^ mice induced P3.5. Boxes in A and B are shown magnified in A’ and B’, respectively. (C) Western blot of Klf4 in cultured dental pulp cells from P7.5 *Bmpr1*α ^*fl/fl*^ control mice treated with Bmp2 or Bmp4 or mock-treated (control). (D) Co-immunoprecipitation experiment using Flag-tagged Smad1 and HA-tagged Klf4 expressed in 293T cells. Smad1 was immunoprecipitated (IP) and immunoblotted (IB) for associated Klf4. (E-E’) Klf4 immunofluorescence after treatment of dissociated apical pulp culture for 48 hours with Ad-m-Klf4 (E) or ad-GFP (E’). (F) qPCR for *Dspp* in explant cultures treated with Ad-m-Klf4 (blue bar) compared with Ad-m-GFP (gray bar). N=3. *, p<0.05. Scale bars, 50μm.

## DISCUSSION

In this work we have identified the MSC population in mouse molars as Gli1+ cells, using *in vivo* genetic cell lineage study and *in vitro* differentiation criteria. Interestingly, BMP signaling is activated when dental mesenchyme cells become committed into odontoblasts, which are adjacent to but exclusive from the most apical Gli1+ MSCs. In addition, we identified potential downstream targets of BMP signaling in root development associated with differentiation defects, including specific transcription factors such as Klf4, Satb2, and Lhx8. Specifically, Klf4 may be a key player in activating the odontogenic mesenchymal lineage commitment. In addition, Pax9 is located in the most apical region overlapping with Gli1+ cells and is upregulated in BMP signaling mutant roots that contain an increased number of uncommitted cells, suggesting Pax9 may be a new marker for MSCs and may help to maintain MSCs in an undifferentiated state. Moreover, our results indicate that BMP signaling activity is required for the lineage commitment of MSCs and, consequently, the initiation of apical growth of the molar root.

### Dynamic activation pattern of BMP signaling may define the stem cell niche environment supporting tooth root growth

We reported here that Gli1+ MSCs are present transiently in the developing molar, and previous studies have shown that they are not detectable in adult molars when tissue growth has ceased (Zhao et al., 2014), consistent with the physiological requirement for finite growth of the molar. The transient nature of these Gli1+ MSCs in the molar root region indicates that they do not maintain self-renewal capacity *in vivo*, one of the defining characteristics that distinguish stem cells from more committed progenitor cells. In contrast, under inductive conditions *in vitro*, both these Gli1+ cells and SCAPs, their putative human counterparts, exhibit the same differentiation potential as adult MSCs as well as show clonogenic ability indicating a self-renewal capability (Sonoyama et al., 2006; Sonoyama et al., 2008). Stem cells have historically been classified as either embryonic stem cells, which are derived from blastocysts, or adult stem cells, which are undifferentiated cells in the body necessary to replenish tissues as they undergo turnover or injury repair. This classification leads to ambiguity in distinguishing cells with stem cell properties from multipotent embryonic precursor cells supporting tissue growth. Increasingly more cell populations with stem cell properties have been identified from multipotent embryonic precursor cells. For example, in the neural crest, a transient multipotent stem cell population was reported to possess high self-renewal ability *in vitro* (Stemple and Anderson, 1992). The transient nature of stem cells utilized during development may be due to a loss of niche environment rather than an intrinsic inability to undergo self-replication as seen in classical adult stem cells. Therefore, it is crucial to gain a better understanding of the stem cell niche environment in order to investigate its functions in controlling the fate of MSCs during tissue growth, homeostasis, and regeneration.

During molar root formation, the zone of BMP activity associated with more committed cells gradually expands as Gli1+ cells disappear, suggesting that the dynamic spatial and temporal regulation of BMP activity may play a role in regulating the stem cell niche disappearance. In contrast, the pattern of BMP activity in the incisors is maintained in a restricted region from newborn stage onwards, adjacent to the incisor Gli1+ MSCs that support the continuous apical growth of the incisor (see Fig. S1). This dynamic niche remodeling adapted for developmental growth supports the notion that these Gli1+ stem cells in molars are *bona fide* stem cells and that the different destinies of Gli1+ stem cells in incisors and molars may be largely associated with changes in the *in vivo* niche environment that promote maintenance or differentiation of stem cells. Increasing evidence supports the idea that niches are not static, but rather persist for varying lengths of time and may be established or disappear at different time points, presumably in response to changing needs of the tissue (Calvi and Link, 2015). Similarly, the niche that harbors the stem cell population supporting tissue growth may gradually disappear as the organism reaches maturity, while a new niche is established to support a population of adult stem cells for tissue repair following injury. For example, although Gli1+ MSCs disappear during root development, their derivatives contribute to the entire pulp tissue including the perivascular cells. This finding suggests that MSCs that support tissue development may be the source of the tissue-specific MSCs recruited to and maintained by the perivascular stem cell niche that function as adult stem cells and participate in injury repair (Bianco et al., 2008; Feng et al., 2011). Taken together, our results demonstrate that murine tooth development offers an excellent *in vivo* model for studying mesenchymal stem cells.

### BMP-regulated transcription factor network controls odontoblast differentiation

Bmps can trigger MSCs to activate transcription factors that are specific to their tissue origin *in vivo*, such as PPARy and Runx2, in adipogenic and osteogenic specification, respectively (Beederman et al., 2013; Wei et al., 2013). Similarly, BMP signaling activates dental pulp MSCs to express odontoblast markers (Iohara et al., 2004; Casagrande et al., 2010). Using our *in vivo* root development model, we found for the first time that in dental pulp cells, Bmps regulates different downstream targets and leads to at least three distinct cell populations, undifferentiated, pulp and odontogenic cells. We have identified transcription factors, Pax9, Klf4, Satb2, Lhx8, and Foxf2, putative mediators of downstream events leading to specific cell fates activated by BMP signaling that are associated with commitment of dental mesenchyme tissue at postnatal stages. These transcription factors are expressed in a spatially restricted pattern consistent with their involvement in determining distinct cellular fates within the dental mesenchyme. Specifically, loss of BMP signaling leads to an altered pattern of this network, suggesting that BMP signaling fine-tunes the spatial distribution of this network, for example, by restricting Pax9 expression and promoting Klf4 expression. Both Pax9 and Klf4 are transcription factors involved in multiple developmental events in a context dependent manner. The tissue specific role of these transcription factors in odontogenesis may be determined by a unique network of transcription factors, in which Pax9 and Klf4 may interact with other members to determine the fate of MSCs during root development.

### BMP-regulated MSCs may be useful for tooth regeneration approaches

The root structure is critical for the ability of the tooth to perform its biological function of occlusion, and clinical treatment of root defects is challenging. Although titanium implants have been used to replace biological roots, the metal-bone osteointegration lacks the ability to respond to the occlusion force. Therefore, an approach utilizing biological tooth root replacement is desirable in clinical dentistry. Human SCAPs have been investigated for their potential use in tooth regeneration approaches (Sonoyama et al., 2006). The apical molar mesenchyme in adult mice does not appear histologically to contain a cell population that resembles the human apical papilla that harbors SCAPs. However, we have identified a cell population, Gli1+ cells, similar to SCAPs, which exhibits MSC characteristics and is capable of supporting tissue growth *in vivo*. We propose that these Gli1+ MSCs may be useful for tooth root regeneration studies in animal models because they meet the necessary criteria of robustly supporting growth and expansion potential. Biological root formation requires a system complex consisting of the interactions between the dental pulp, periodontium, and epithelial cells. To mimic this biological process for tooth regeneration, future studies will need to investigate the coordination between these components.

In addition, previous studies have shown that epithelial-mesenchymal interactions are critical for tooth development as well as many other developmental processes (Li et al., 2017). At early embryonic stages, BMP signaling is relayed between the epithelium and mesenchyme as tooth development progresses. At newborn stage, BMP signaling becomes specifically required in the mesenchyme but not in the epithelium when root formation begins after crown development is completed. BMP signaling regulates cell-specific downstream events in a context dependent manner and generally directs stem cells to undergo differentiation according to their tissue origins, possibly through activation of lineage-specific transcription factors. We note that the BMP pathway may be a good access point for manipulating multiple pathways and identifying downstream targets critical for cell fate determination. Moreover, BMP regulatory mechanisms that we have begun to dissect in detail may help guide future tooth regeneration approaches.

## MATERIALS AND METHODS

### Generation of transgenic mice

The *Gli1-CreER* knock-in (JAX#007913, The Jackson Laboratory, Ahn and Joyner, 2004), *tdTomato* conditional reporter (JAX#007905, The Jackson Laboratory, Madisen et al., 2010), conditional *Bmpr1a* floxed (Gift from Sarah E. Millar, University of Pennsylvania, Andl et al., 2004), *K14-rtTA* (JAX#007678, The Jackson Laboratory, Xie et al., 1999), *tetO-Cre* (JAX#006234, The Jackson Laboratory), and *Gli1-LacZ* knock-in/knock-out reporter (JAX#008211, The Jackson Laboratory, Bai et al., 2002) mouse lines have all been described previously. *Gli1-LacZ* knock-in/knock-out mice were used as heterozygotes.

### Tamoxifen and doxycycline administration

Tamoxifen (Sigma T5648) was dissolved in corn oil (Sigma C8267) at 20 mg/ml and injected intraperitoneally (i.p.) at a dosage of 1.5mg/10g body weight. Doxycycline rodent diet (Harlan, TD.01306) was administered every day.

### X-gal staining and detection of β-galactosidase activity

Samples at various stages of postnatal development were fixed in 0.2% glutaraldehyde, decalcified in 10% ethylenediaminetetraacetic acid (EDTA, pH7.4) passed through a sucrose series, embedded in OCT Compound (Tissue-Tek) and sectioned on a cryostat at 10 μm prior to X-gal staining for lacZ expression. To detect of β-galactosidase (β-gal) activity in tissue sections, cryosections were stained in X-gal staining solution (2mM MgCl_2_, 0.01% sodium deoxycholate, 0.005% Nonidet P-40, 5mM potassium ferricyanide, 5mM potassium ferrocyanide, 20mM Tris pH7.3, and 1 mg/ml X-gal in PBS) for 3-4 hours at 37°C in the dark, followed by postfixation in 4% paraformaldehyde for 10 min at room temperature and counterstaining with nuclear fast red (Electron Microscopy Sciences, 2621203).

### Histological analysis

Dissected samples were fixed in 4% paraformaldehyde overnight at 4°C and then decalcified in 10% DEPC-treated EDTA (pH 7.4) for 1-4 weeks depending on the age of the sample. Samples were passed through serial concentrations of ethanol for embedding in paraffin and sectioned at 7μm thickness using a microtome (Leica). Deparaffinized sections were stained with Hematoxylin and Eosin (H&E) using standard procedures for general morphology.

For cryosections, decalcified samples were dehydrated in 60% sucrose/PBS solution overnight at room temperature. Samples were then embedded in OCT compound (Tissue-Tek, Sakura) and frozen onto a dry ice block to solidify. Embedded samples were cryosectioned at 7μm thickness using a cryostat (Leica CM1850).

### In situ hybridization

After deparaffinization and serial hydration with ethanol, sections were treated with proteinase K (20 mg/ml) for 5 min at room temperature and post-fixed in 4% paraformaldehyde in PBS for 10 min. After treatment in 0.1M triethanolamine with 0.25% acetic anhydride for 10 min, sections were dehydrated through serial concentrations of ethanol and air-dried. Digoxigenin labeled probes in hybridization solution were heated in a 100°C water bath for 5 min, chilled on ice and incubated with the slides in the humidified hybridization chamber at 65°C for 16 hours. After hybridization, sections were treated with 1 µg/ml RNase A (Sigma) in 2X SSC buffer for 30 min at 37°C, and washed in 3 times in prewarmed 2X SSC buffer and 0.2X SSC buffer with 0.05% CHAPS (20 min each) at 65°C. The sections were blocked with 20% sheep serum/PBST for 2 hours at room temperature and incubated with 1:5000 dilution of anti-digoxigenin-ap antibody (Roche, 11093274910) at 4°C overnight. After overnight incubation, sections were washed 3 times with PBS with 0.1% Tween 20 and 1mM tetramisole hydrochloride for 10 min each, followed by a wash with alkaline-phosphatase buffer (100mM NaCl, 100mM Tris-HCl pH9.5, 50mM MgCl_2_, and 0.1% Tween 20 in H_2_O). Finally, sections were developed using the BMPurple substrate system (Roche). *Dspp* cDNA clones were kindly provided by Irma Thesleff (University of Helsinki, Finland). *Foxf2 and Lhx8* digoxigenin-labeled anti-sense RNA probes were generated using *Foxf2* cDNA (NCBI Reference: NM_010225.2, 1084-1955) and *Lhx8* cDNA (NCBI Reference: NM_010713.2, 541bp-1608bp) as templates, respectively.

### Immunostaining

Sections were immersed in preheated antigen unmasking solution (Vector, H-3300) in an Electric Pressure Cooker (976L, Cell Marque, Sigma) at high pressure for 10 minutes, followed by cooling at room temperature for 30-45 min and incubation with blocking reagent (PerkinElmer, FP1012) for 1 hour and then primary antibody overnight at 4°C. After 3 washes in PBS, sections were incubated with Alexa-conjugated secondary antibody (Invitrogen). For pSmad1/5/8, bHRP-labeled goat anti-rabbit IgG (PerkinElmer, NEF812001EA; 1:200) was used as secondary antibody and TSA kits were used for signal detection (PerkinElmer, NEL741001KT). Sections were counterstained with DAPI (Sigma, D9542). Images were captured using a fluorescence microscope (Leica DMI 3000B) with filter settings for DAPI/FITC/TRITC.

Immunostaining was performed using the following antibodies: pSmad1/5/8 (1:500, Cell Signaling, #9511), Ki67 (1:100, Abcam, ab16667), β-gal (1:50, Abcam, ab9361), Pax9 (1:25, Abcam, ab28538), Satb2 (1:100, Abcam, ab92446), and Klf4 (1:25, Sigma, HPA002926). Alexa Fluor 568 and Alexa Fluor 488 (1:200, Invitrogen) were used for detection.

### Clonal culture and multipotential differentiation

The apical one third of the dental pulp mesenchyme from the mandibular molar region of P3.5 mice was separated, minced, and digested with solution containing 2 mg/mL collagenase type I (Worthington Biochemical, New Jersey, USA) and 4 mg/mL dispase II (Roche Diagnostics, California, USA) in PBS for 1 hour at 37°C. A single-cell suspension was obtained by passing the cells through a 70-µm strainer (BD Biosciences, California, USA) and was seeded at 1 × 10^5^/well in 6-well plate culture dishes (Corning, New York, USA) with α-MEM supplemented with 20% FBS, 2 mM L-glutamine, 55 µM 2-mercaptoethanol, 100 U mL-1 penicillin, and 100 µg/mL streptomycin (Life Science Technologies). The culture medium was changed after an initial incubation for 48 hours and the attached cells were cultured for another 14 days at 37°C under hypoxic conditions (5% O_2_, 5% CO_2_, balanced with nitrogen). Clones could be detected 7-10 days after plating.

For the differentiation assays, cells from the colonies were cultured until confluent and then induced in osteogenic, adipogenic or chondrogenic differentiation medium (05504, 05503, 05455, Stemcell Technologies, Vancouver, Canada) for 2-3 weeks according to the manufacturer’s instructions.

### RNA sequencing (RNA-seq) analysis

*Gli1-CreER;Bmpr1*α ^*fl/fl*^ and *Bmpr1*α ^*fl/fl*^ littermate control mice received tamoxifen at P3.5 days and were euthanized 4 days thereafter. The apical half of the first mandibular molars was dissected out and RNA was extracted using the RNeasy Plus Mini Kit (Cat. 74134, Qiagen). cDNA library preparation and sequencing were performed at the Epigenome Center of the University of Southern California. A total of 200 million single-end reads were obtained on Illumina NextSeq500 equipment for 3 pairs of samples. High quality reads were aligned to mm10 using TopHat 2 in conjunction with a gene model from Ensembl release 61. Data was quantitated by counting the number of reads over exons and normalized as RPKM (reads per kilobase per million mapped reads) (Mortazavi et al., 2008). The values were adjusted globally by matching count distributions at the 75^th^ percentile and then adjusting counts to a uniform distribution between samples. Differential expression was estimated by selecting transcripts that displayed significant changes (p < 0.05) after Benjamini and Hochberg correction using a null model constructed from 1% of transcripts showing the closest average level of observation to estimate experimental noise as detailed previously (Kim et al., 2016). The gene list was further ranked using fold change criteria. For visualization, aligned reads were uploaded to Integrated Genome Viewer (IGV, Broad Institute). The raw data has been deposited to NCBI under the accession number GSE79791. To investigate potential functional enrichment of various biological pathways in differentially expressed genes in RNA-seq, a ranked p-value was computed for each pathway from the Fisher exact test based on the binomial distribution and independence for probability of any gene belonging to any enriched set as detailed previously (Kim et al., 2016).

### Western blot and co-immunoprecipitation

Dental pulp apical cells from the apical region of P3.5 molar were harvested 48 hours after treatment with 10ng/ml Bmp2 (355-BM-010, R&D systems) or Bmp4 (5020-BP-010, R&D systems), or mock treated with equal volume of solvent (0.1% bovine serum albumin in 4 mM HCl) as control. For immunoblotting, cells were lysed in lysis buffer (50 mM Tris-HCl pH 7.5, 150 mM NaCl, 2mM EDTA, 0.1% NP-40, 10% glycerol, and protease inhibitor cocktail). After protein quantification using Bio-Rad protein assays (Bio-Rad Laboratories), 20-80 μg of protein was separated by SDS-PAGE and transferred to 0.45 μm PVDF membrane. Membranes were blocked in TBS with 0.1% Tween 20 and 5% BSA (blocking solution) for 1 hour, followed by overnight incubation with primary antibody diluted at 1:500-1:5000 in blocking solution, and 1 hour incubation with HRP-conjugated secondary antibody diluted at 1:5,000–1:20,000. Immunoreactive protein was detected using ECL (GE Healthcare) and BioMax film (Kodak) or FluorChem E Chemiluminescent Western Blot Imaging System (Cell Biosciences).

For immunoprecipitation, HA-Klf4 (Plasmid #34593, Adgene) and Flag-Smad1 (Alliston et al., 2005) plasmids were transfected into 293T cells. Cells were harvested 24 or 48 hours after transfection and lysed in lysis buffer (50 mM Tris-HCl pH 7.5, 150 mM NaCl, 2mM EDTA, 0.1% NP-40, 10% glycerol, and protease inhibitor cocktail). Lysates were subjected to immunoprecipitation with anti-Flag antibody and protein G-Sepharose 4 fast flow (GE Healthcare). Immune complexes were washed three times with lysis buffer and subjected to immunoblotting with anti-Flag (Sigma, F1804) or anti-Klf4 antibodies.

### Adenovirus treatment

GFP control adenovirus (Ad-GFP, Cat. 1060) and Klf4 overexpression adenovirus (Ad-m-KLF4, 1791) were purchased from Vector Biolabs. To validate the Klf4 expression after virus infection, Ad-m-KLF4 adenovirus was added to the dental pulp cell culture for 3 hours. 48 hours later, the cells were fixed in 4% PFA on ice for 10 min and immunostained with Klf4 (1:100). For functional analysis of Klf4, the apical region of P5.5 mouse molars was cut into small pieces to facilitate virus penetration. Ad-m-KLF4 adenovirus was added to the dissociated apical pulp culture and cultured for 48 hours. Five days later, RNA was extracted from cultured pulp tissues using RNeasy Micro Kit (74004, Qiagen) and reverse-transcribed to cDNA (Sensiscript, 205211, Qiagen) for qPCR analysis using Dspp qPCR primers (PPM40292F, Qiagen).

### Sample size and statistics

N=3 for all experiments unless otherwise stated. Student’s t test was applied for statistical analysis. A p-value of <0.05 was considered statistically significant.

## ACKNOWLEDGEMENTS

We thank J. Mayo for critical reading of the manuscript. We thank Epigenome Center of the University of Southern California (USC) for performing RNA sequencing. We thank the Norris Medical Library Bioinformatics Service, funded by the USC Office of Research and the Norris Medical Library, for assisting with sequencing data analysis.

## Author contributions

J.F. and Y.C. designed the study and wrote the manuscript. J.F. performed the experiments and analyzed the data. J.J., J.L., H.Z. participated in transgenic animal experiments. T.Z. and J.X. participated in the biochemistry experiments. V.P. performed the bioinformatics analysis for the RNA sequencing data.

## Data availability

RNA sequencing data has been submitted to NCBI. It has been assigned #GSE79791

## Supplementary Table and Figure Legends

**Table S1.**
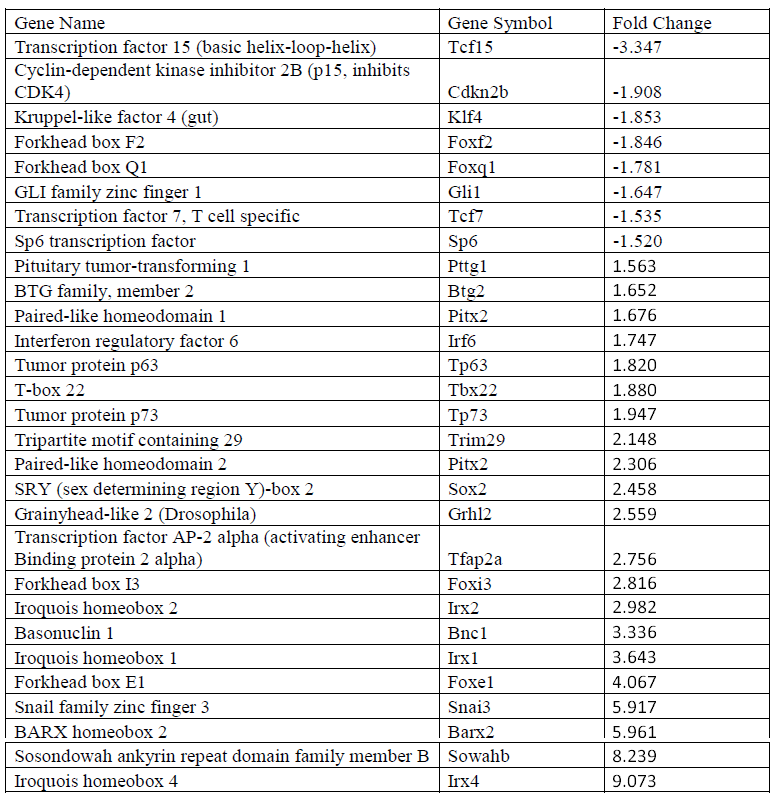
Differentially expressed transcription factors in the apical region of *Gli1-CreER;Bmpr1* α^*fl/fl*^ molars compared with *Bmpr1*α^*fl/fl*^ control molars Down-and up-regulated genes identified by RNA-seq of the apical half of the molar mesenchyme from *Gli1-CreER;Bmpr1*α^*fl/fl*^ mice compared with Bmpr*1*α^*fl/fl*^ control mice at P7.5 after induction at P3.5. (Fold change >1.5, p<0.05)

**Figure S1. Activated BMP signaling in molars and incisors at different stages**. (A-D) pSmad1/5/8 (red) immunostaining of E18.5 (frontal section; A) and P3.5, P7.5 and P13.5 (sagittal sections; B-D) sections of mandibular molars from C57/BL6 wild type mice. (E-H) pSmad1/5/8 (red) immunostaining of sagittal sections of mandibular incisors from P3.5 and 6-week-old adult C57/BL6 wild type mice. Scale bars, 100μm.

**Figure S2. Postnatal epithelial BMP signaling is dispensable for root development**. H&E staining of sagittal sections of mandibular molars of P21.5 littermate *Bmpr1*α^*fl/fl*^ control (A-C) and *K14-rtTA;tetO-Cre;Bmpr1*α^*fl/fl*^ (D-F) mice after doxycycline induction from P6.5 to P21.5. B, C and E, F are magnified views of distal and proximal roots from A and D, respectively. Scale bars, 100μm.

